# Human stem cell models for group 3 medulloblastoma uncover JARID1B as a regulator of the chromatin landscape

**DOI:** 10.64898/2025.12.06.689939

**Authors:** Zulekha A. Qadeer, Elise Hou, Dara Siegel, Samantha Westelman, Brian Gudenas, Kyle Smith, Wazim Mohammed Ismael, Linyu Wang, Mackenzie O Johnson, Shane Grele, Ronald Phua, Liam Hendrikse, Ekin Guney, Miller Huang, Alexandre Gaspar-Maia, Paul A. Northcott, Michael D. Taylor, William A. Weiss

## Abstract

Medulloblastoma (MB) is one of the most prevalent malignant brain tumors in children, with tremendous cognitive and neuroendocrine disability among survivors. Group 3 MB (G3MB) has poor overall survival at <50%, high frequencies of metastases, and no targeted therapies. Amplification of *MYC* and activation of TGFβ signaling occur frequently in G3MB. Many tumors have no reported mutations, suggesting epigenetic drivers. We here describe novel humanized models for G3MB from human induced pluripotent stem cells (hiPSC). By transducing hiPSC-derived neuroepithelial stem cells (NESC), we determined that: 1) both MYC and TGFβ effectors drove tumors *in vivo*; 2) MYC/TGFβR1 in combination led to more aggressive tumors and resistance to clinical inhibitors of TGFβ, and 3) NESC-derived tumors clustered with human G3MB. To decipher mechanisms, we integrated RNA-sequencing with CUT&RUN (for MYC genomic localization and post-translational modification of histones). MYC-bound neural developmental genes were repressed in MYC/TGFβR1 co-driven lines. Gene signatures associated with the Polycomb Repressive Complex (PRC) demarcated with H3K27me3; the histone mark directly regulated by PRC. We identified JARID1B, a MYC binding partner and H3K4me3 demethylase, as a regulator of repressed neural genes. Primary G3MB also showed increased levels of H3K27me3 concurrent with higher expression of JARID1B. Knockdown of JARID1B in human G3MB cell lines reduced growth, supporting potential as a therapeutic target. We conclude that a MYC-TGFβ-JARID1B axis represses target genes to drive G3MB and present new humanized models for G3MB to understand epigenetic dysregulation in G3MB.

## Introduction

Medulloblastoma (MB) is the most common brain tumor in children and typically develops in the posterior fossa. Most survivors experience long-term sequelae caused by highly aggressive multimodal therapies^1–3^. There is an imperative to identify new approaches that target tumor over normal brain, sparing cognition and development. Genomic studies have revealed 4 genetically and epigenetically distinct subgroups (WNT, SHH, Group 3, Group 4)^4–7^. Children with Group 3 MB (G3MB, 20% of patients) are much more likely to succumb to their disease. Moreover, G3MBs have few recurrent mutations, high frequencies of metastasis to other parts of the central nervous system (CNS) and no targeted therapies^3,7^.

Genomic analyses have provided extensive information about genes aberrantly expressed or mutated in MB and defined key subgroup drivers. Notably, *MYC* amplification is common in G3MB. TGFβ effectors are also activated frequently through copy number variations or gain of enhancers ^8,9^. The TGFβ signaling pathway has pleiotropic functions regulating cell growth, differentiation, migration, invasion, angiogenesis, and immune regulation. TGFβ pathway deregulation has crucial roles in tumor initiation, development, and metastasis, that can be tumor suppressive or promoting depending on context^10,11^. This complexity is further challenged by intratumoral heterogeneity mediated by TGFβ, which poses a challenge to cancer treatment^12^. The functional significance of MYC and the TGFβ pathway in G3MB is currently not well defined.

Chromatin dysregulation activates and maintains aberrant transcriptional programs in pediatric cancers. DNA methylation, chromatin modification and post-translational modifications (PTMs) of histones play major roles in chromatin-regulated processes and transcriptional readouts^6^ ^13^. Histone PTMs in particular modulate interactions between DNA and histones at *cis-*regulatory elements to impact gene regulation. Such crosstalk at enhancers and promoters culminates in transcriptional diversity of genetically uniform individual cells within a tumor^9,14–16^. Unlike WNT and SHH subgroups, where alterations frequently activate those pathways, ∼30% of genes deregulated in G3 tumors converge on epigenetic modifiers and regulators of transcription^15,17,18^. Moreover, epigenetic and transcriptional states render human fetal rhombic lip progenitors vulnerable to transforming to this MB subtype^17,19^. These states are annotated by the interplay between Polycomb Repressive Complex (PRC) and the MLL/COMPASS family^20–22^. MLL/COMPASS enzymes establish the H3K4me3 mark associated with gene activation. PRC2 is a histone methyltransferase with distinct catalytic functions in chromatin regulation generally associated with transcriptional silencing via deposition of H3K27me3^21^. Inactivating mutations in MLL/COMPASS family members *MLL2/KMT2D*, *MLL3/KMT2C*, and *KDM6A/UTX* occur in 16% of MB^15^ and repress neural differentiation genes in SHHMB^23^. How these mutations alter COMPASS formation, cooperate with MYC amplification and engage with PRC2 and the H3K27me3 mark in G3MB remains unexplored. Importantly, an outstanding question is how these complexes associate during human cerebellar development and how crosstalk on the chromatin regulates the rhombic lip-lineage trajectory.

During development, embryonic stem cells (ESC) differentiate into three germ layers. As neural ectoderm forms the neural tube, neural stem cells (NSC) are born in the neuroepithelium and further differentiate into granule neuron precursors (GNPs) in the external granule layer that populate the cerebellum in the hindbrain, the primary site of MB^24,25^. Distinct cellular origin and developmental pathways are the likely basis for the striking clinical heterogeneity of MB and specifically impact clinical outcomes^26^. The developing human cerebellum has anatomical features that are unlike those of most other mammals. In particular, the human rhombic lip (RL), which gives rise to all glutamatergic neurons, is divided into a relatively primitive and quiescent ventricular zone (RL^VZ^) and a more differentiated and proliferative subventricular zone (RL^SVZ^)^24,25^. G3MB arises from an early RL^SVZ^ lineage^17,19^. To faithfully recapitulate G3MB, human stem cell-based models are therefore needed.

The use of iPSC-derived human NESC culture systems advances our understanding of human brain development and disease by providing a manipulatable and scalable cell population for biochemical and genetic studies not feasible in human brain^27,28^. Further, human cells hold promise to recapitulate the genetic and epigenetic architecture of human cancer more accurately compared to rodent cells. We used hiPSCs generated via integration free methods and derived NESCs. Amplification of *MYCN* is found in high-risk SHHMB. We have established that *MYCN* can drive MB in genetically engineered mouse models (GEMM)^29^ and hiPSC-derived NESCs^29^. Given the unique developmental context of G3MB, we sought out to develop humanized models using the hiPSC platform. We hypothesize that MYC and TGFβ pathway represents critical driving events in G3MB and therefore would be excellent candidate genes to utilize to develop these models. In this study, we describe a new humanized model for G3MB driven by MYC and TGFβ pathway effectors and reveal an unexpected role for MYC in gene repression that can help identify new therapeutic vulnerabilities.

## Results

### Human stem cells transduced with MYC and TGFβ pathway effectors model G3MB

We derived NESCs from the TP53^WT^ hiPSC line WTC10^29^ and transduced them with TGFβ pathway effectors activated in MB (ACVR2A, TGFβR1, and TGFβ1)^8,9^ either alone and/or in combination with MYC, prioritizing combinations observed in patients (**Fig. S1A-B**). Overexpression of MYC and TGFβ pathway effectors alone or in combination led to increased proliferation compared to control NESCs, with the most significant acceleration in the combined MYC and TGFβ pathway cell lines (**Fig. S1C**). MYC contributes to therapy resistance across many cancer types^30^. We therefore treated NESCs with galunisertib (LY2157299) and LY3200882^10,31^, clinical small molecule inhibitors that target TGFβR1. Both drugs significantly reduced proliferation over time in NESCs expressing TGFβ1 and TGFβR1. This phenotype was ablated in cells overexpressing MYC and TGFβ effectors in combination. Of note, we didn’t see any proliferation differences in control NESCs, indicating specificity of these inhibitors to pathway overexpression driven by TGFβR1 (**Fig. S1D**). We also treated MYC and TGFβR1expressing NESCs with cyclophosphamide and carboplatin, standard of care agents for patients with MB. TGFβR1 alone and MYC combined with TGFβR1 promoted resistance to both agents (**Fig. S1E**). These collective data support that the combination of MYC and TGFβ promotes therapy resistance to various agents in G3MB.

We next implanted NESCs orthotopically into immunocompromised (NSG) mice. Individually, MYC, TGFβ1 effectors each drove tumor growth (**Fig. 1B-C, Fig S2A**). When combined with MYC, TGFβ pathway effectors (ACVR2A, TGFβ1 and TGFβR1) accelerated tumorigenesis compared to MYC or each TGFβ effector alone. We also observed leptomeningeal spreading and metastases to the spine of mice injected with MYC-TGFβR1 NESCs in the lateral ventricles at an accelerated rate, compared to MYC NESCs (**Fig. 1D**). Histological analysis of NESC-tumors confirmed features typical of human MB including small, rounded nuclei with prominent nucleoli, Homer Wright rosettes, mitotic figures, and necrosis. Tumors derived from MYC-TGFβ effectors were the most aggressive and less differentiated with highly disorganized chromatin and oversized nucleoli (**Fig. S2B**). We performed RNA-seq analysis on NESC-derived tumor and found they clustered to G3/G4MB tumors (**Fig. 1E**). These tumors also had higher expression of G3MB driver genes OTX2^8,19^ and HLX^14^ (**Fig. 1F, Table S1**). Collectively, we validated that MYC, TGFβ and MYC/TGFβ stem cell models as isogenic systems for G3MB, and align with basic science observations that OTX2 is a critical target of TGFβ signaling in the developing nervous system^32^.

**Figure 1.**
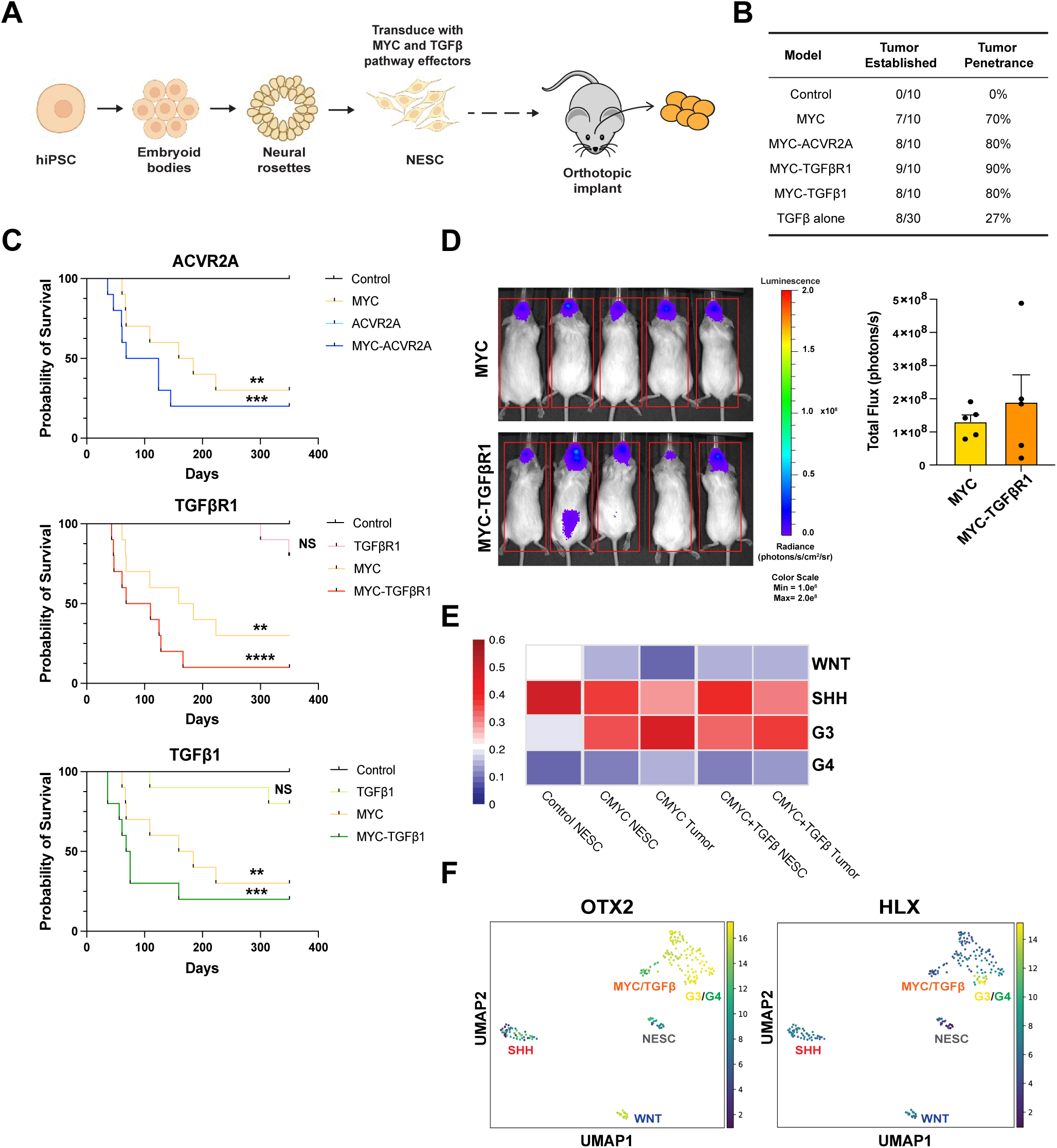
MYC and TGFβ effectors promote group 3 medulloblastoma *in vivo*. **A,** Kaplan Meier survival curve of mice orthotopically implanted with NES cells transduced as described in Fig. 1A (n=5). **B,** Table summarizing tumor penetrance for each genotype. **C,** Representative bioluminescent imaging data from n=5 mice orthotopically injected with MYC and MYC-TGFβR1 NESCs, measured in relative light units **D,** Canonical correlation analysis of bulk RNA-seq data from hiPSC-derived NESCs and tumors with human MB tumors (Smith et al., 2022). **E,** UMAP plots of principle component analysis of cells and tumor types for select G3MB gene expression.

### MYC-TGFβ gene signatures associate with PRC-interactors and neuronal differentiation

Data in Fig 1 suggest that combined MYC and TGFβ overexpression promote aggressive tumor formation and intrinsic drug resistance. To address mechanisms, we queried RNA-seq of NESC-derived tumors for unique expression signatures *in vivo* (**Fig 2A**). We compared differential expression profiles of tumors driven by MYC alone and TGFβ alone versus MYC-TGFβ pathway effectors (ACVR2A, TGFβ1 and TGFβR1) to identified 597 differentially enriched genes shared among tumors driven by MYC TGFβ pathway effectors (**Fig. 2B, Fig. S3A, Table S1**). Terms from Gene Set Enrichment Analysis (GSEA)^33^ analysis identified chromatin factors enriched MYC-ACVR2A, MYC-TGFβR1, MYC-TGFβ1 tumors (**Fig. S3B**). ChIP-Seq Enrichment analysis (ChEA)^34^ analysis of the MYC-TGFβ gene signature identified these genes as PRC-regulated^35,36^ (**Fig. 2C, Table S1**). Therefore, we focused on identifying important pathways that were upregulated or downregulated exclusively in MYC-TGFβ derived NESC-tumors (**Fig. 2D**). Terms from Gene Ontology (GO)^37^ analysis on the MYC-TGFβ exclusive genes demonstrated neuron channels in the downregulated genes. GO terms enriched in MYC-TGFβ exclusive Up genes were negative regulators of neuron differentiation and wound healing, supporting tumor invasive features (**Fig. 2D-E, Table S1**).

**Figure 2.**
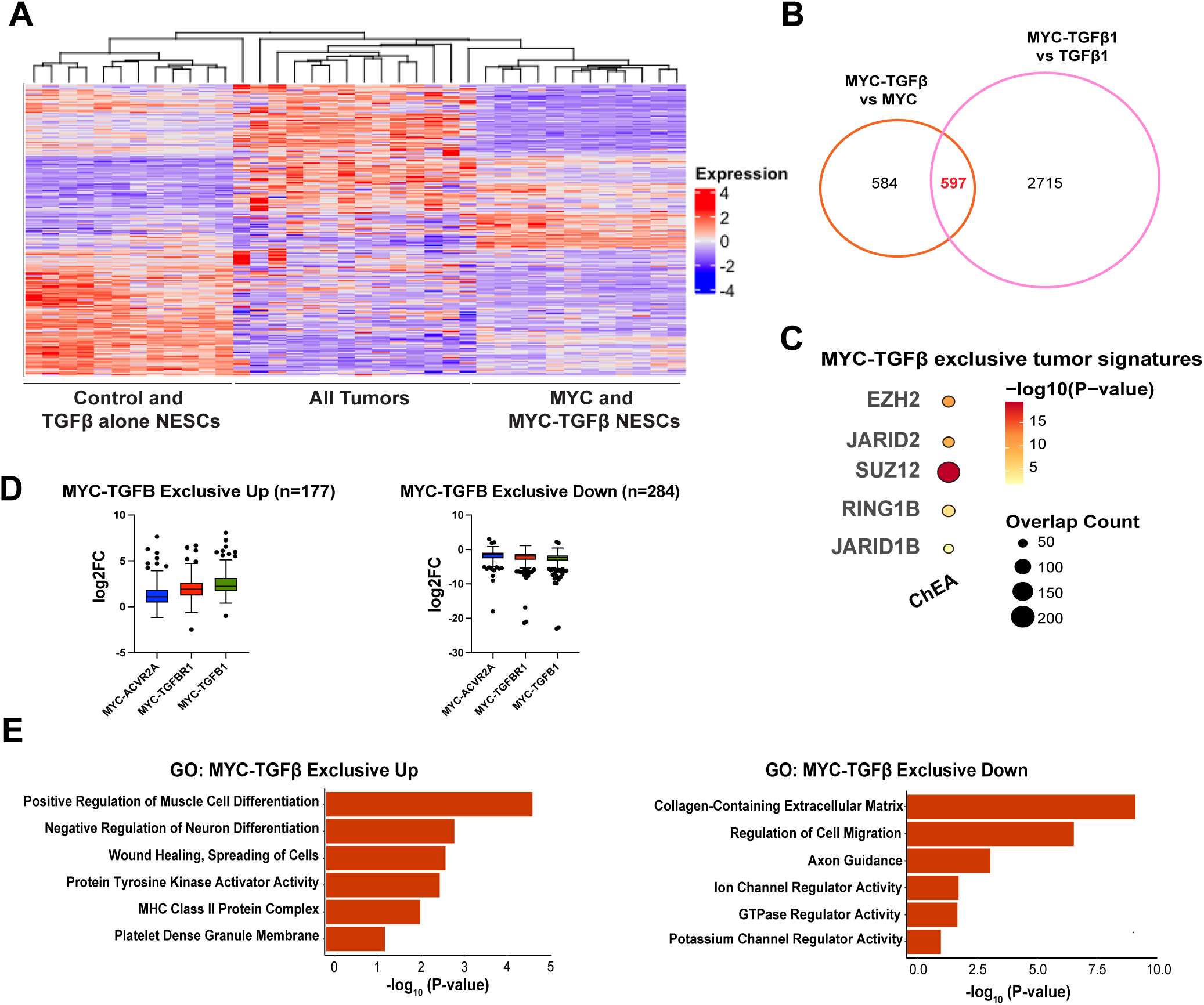
Gene signatures in MYC-TGFβ driven tumors are associated with JARID1B and PRC regulators. **A**, Heatmap of control, MYC, and TGFβ NESCs and tumors shows distinct clusters in MYC alone, TGFβ alone, and MYC-TGFβ tumors. **B**, Venn diagram comparing overlap of differentially significant genes to identify MYCTGFβ exclusive gene signatures in tumors. **C**, Dotplot of significant transcription factors identified in ChIP-sequencing Enrichment Analysis (ChEA) in the common differentially upregulated genes identified in MYC+TGFβ tumors versus MYC alone tumors (n=597). **D**, Boxplot of expression levels of genes upregulated or downregulated exclusively in MYC-TGFβ tumors. **E**, GO analysis of MYC-TGFβ tumors genes from D showing suppression of neural differentiation genes.

### MYC promoter-bound regions are demarcated with gains of H3K27me3 and JARID1B

To identify specific chromatin factors implicated in RNA-seq analyses (Fig S3B), we performed Cleavage Under Targets & Release Using Nuclease (CUT&RUN) for FLAG-tagged MYC genomic localization and activating histone PTMs H3K27ac and H3K4me3 as well as repressive histone mark H3K27me3. Genome wide FLAGMYC enrichment in MYC-NESCs correlated strongly to H3K4me3 and H3K27ac, with significant peaks found primarily in promoters. (**Fig. 3A, Fig. S4A).** Strikingly, FLAG-MYC in MYC-TGFβR1 NESCs enrichment was at regions of H3K27me3, a repressive mark not typically associated with MYC transactivation (**Fig. S4A**). While a significant number of peaks were present in promoters, MYC peaks were also gained in intronic regions and distal regions, suggesting genomic relocalization (**Fig. 3A**). A small subset of significant FLAG-MYC peaks demarcated with H3K27me3 domains were also observed in MYC-TGFβR1 NESCs (**Fig. 3B**). ReMapEnrich^38^ analysis identified significantly enriched regions associated with PRC members and with the H3K4me3 demethylase JARID1B (**Fig. 3C, Table S2**), correlating with the RNA-seq analysis in MYC-TGFβ driven NESC-derived tumors (**Fig. 2C**). Integrating the MYC-TGFβ exclusive Down genes with MYC/H3K27me3 promoter bound genes identified in MYC-TGFβR1 NESCs we identified genes important in extracellular matrix organization, neuron differentiation and synaptic transmission (**Fig. 3D**). Among these genes were critical neurodevelopment transcription factors *PAX5, EN1* and *EN2* and transmembrane proteins *MMP15, MMP25,* and *TENM4* (**Table SX**), supporting a role for MYC-mediated gene repression of neuronal differentiation and impaired extracellular matrix modeling.

**Figure 3.**
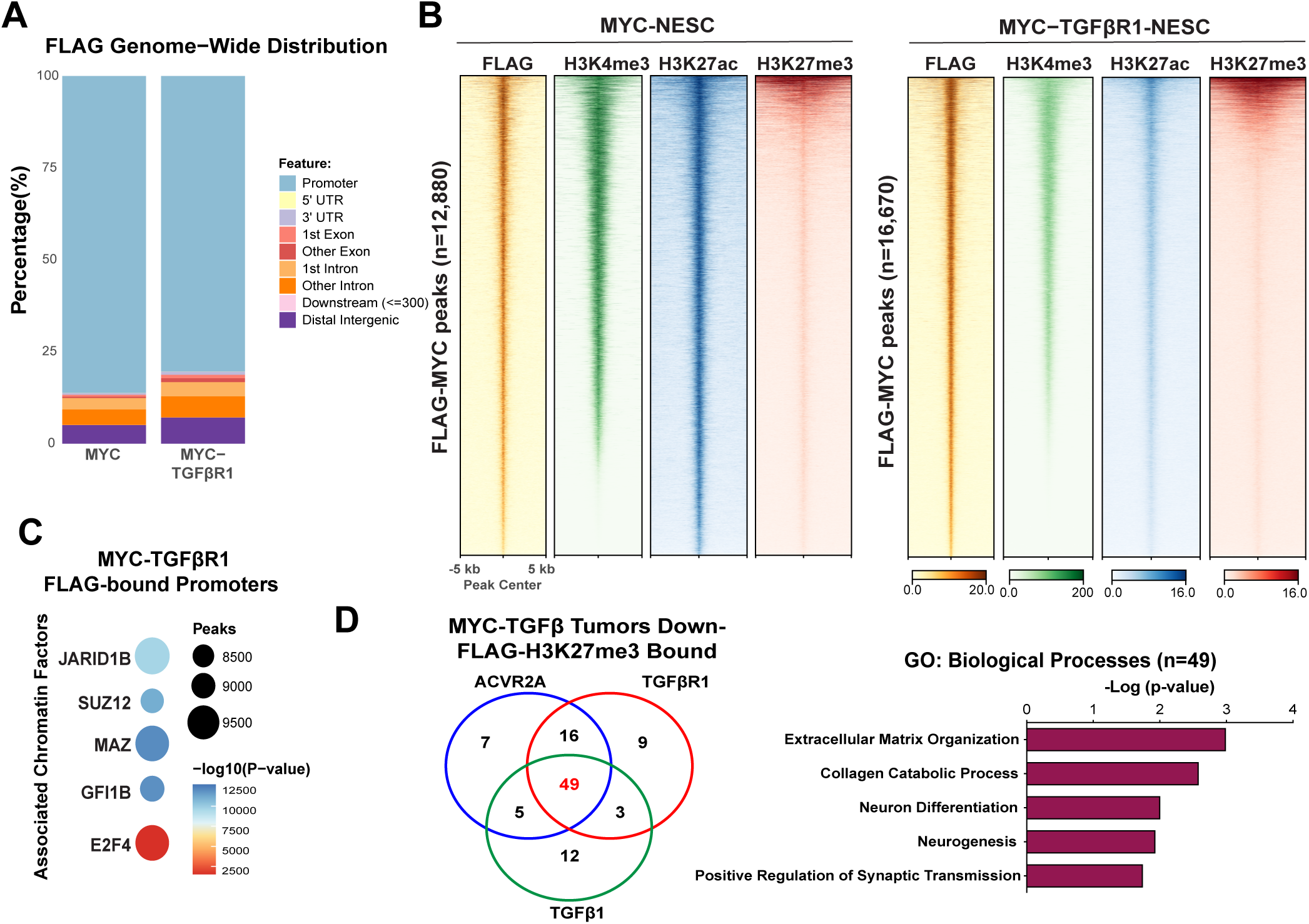
MYC-bound promoters are associated with JARID1B in MYC-TGFβR1 NESC cells. **A**, Genome wide mapping of significant MYC peaks in MYC and MYC-TGFβR1 NESCs. **B**, Heatmap of FLAG-MYC, H3K4me3, H3K27ac and H3K27me3 in MYC and MYC-TGFβR1 NESCs at MYC-bound peaks (±5 kb from peak center). **C**, Motif analysis of MYC-TGFβR1 promoter peaks from A demonstrating number of statistically significant enriched TFs and overlapping genomic regions based off publicly available ChIP-seq databases. Analysis performed using ReMapEnrich. **D**, Venn diagram (left) comparing overlap of MYC-TGFβ exclusive down genes in tumors with MYC-H3K27me3 bound genes in MYC-TGFβR1 cells. GO analysis of MYC-TGFβ down and MYC-H3K27me3 bound (n=49) genes.

Consistent with results in Fig 3, Lee et al recently observed that overall peak numbers and genome coverage of H3K27me3 were significantly higher globally in G3MB compared to SHHMB and G4MB^18^. Given our observations in human G3MB NESC-derived models, we reasoned that PRC deregulation could be suppressing neuronal differentiation through silencing of developmental transcription factors^18^. We therefore performed additional CUT&RUN in human tumor-derived *MYC* amplified G3MB cell lines (**Fig. S4B)**. Analyzing histone PTMs H3K4me3 and H3K27me3 in D283 and D425 cells, we observed broad distribution of H3K27me3 throughout the genome and H3K4me3 peaks primarily at promoters (**Fig. S4C**). We integrated these datasets with control-NESC H3K27me3 enrichment and analyzed differential peak calling with Diffbind^39^ to identify MYCH3K27me3 bound promoters identified in human tumor lines (**Fig. S4D, Table SX)**. Notably, we found correlations of H3K27me3 peaks with JARID1B and MYC regulated regions in D283 and D425 (**Fig. S4E**). These findings suggest that in MYC-driven G3MB, deposition of H3K27me3 converges on a shared set of JARID1B regulated neuronal differentiation genes across both human stem cell-derived models and tumor-derived cells.

### JARID1B is a therapeutic target in human G3MB

The JARID1B (*KDM5B/PLU-1*) histone demethylase catalyzes removal of H3K4me3, silencing H3K4me3 associated genes. JARID1B is recruited by the PRC2 subunit SUZ12 to coordinate silencing of neuronal lineages during mESC differentiation^40,41^. While expression of *JARID1B* is restricted in most normal adult tissues, JARID1B is often upregulated in cancer^42–45^. Since PRC-interactors and JARID1B suppressed signatures of neuronal differentiation (**Fig 2C**, **3C)**, we queried two datasets, observing significant expression of JARID1B with MYC in human G3MB, as well as high levels of EZH2 and SUZ12 in both G3MB and G4MB (**Fig. 4A, S5A**).

**Figure 4.**
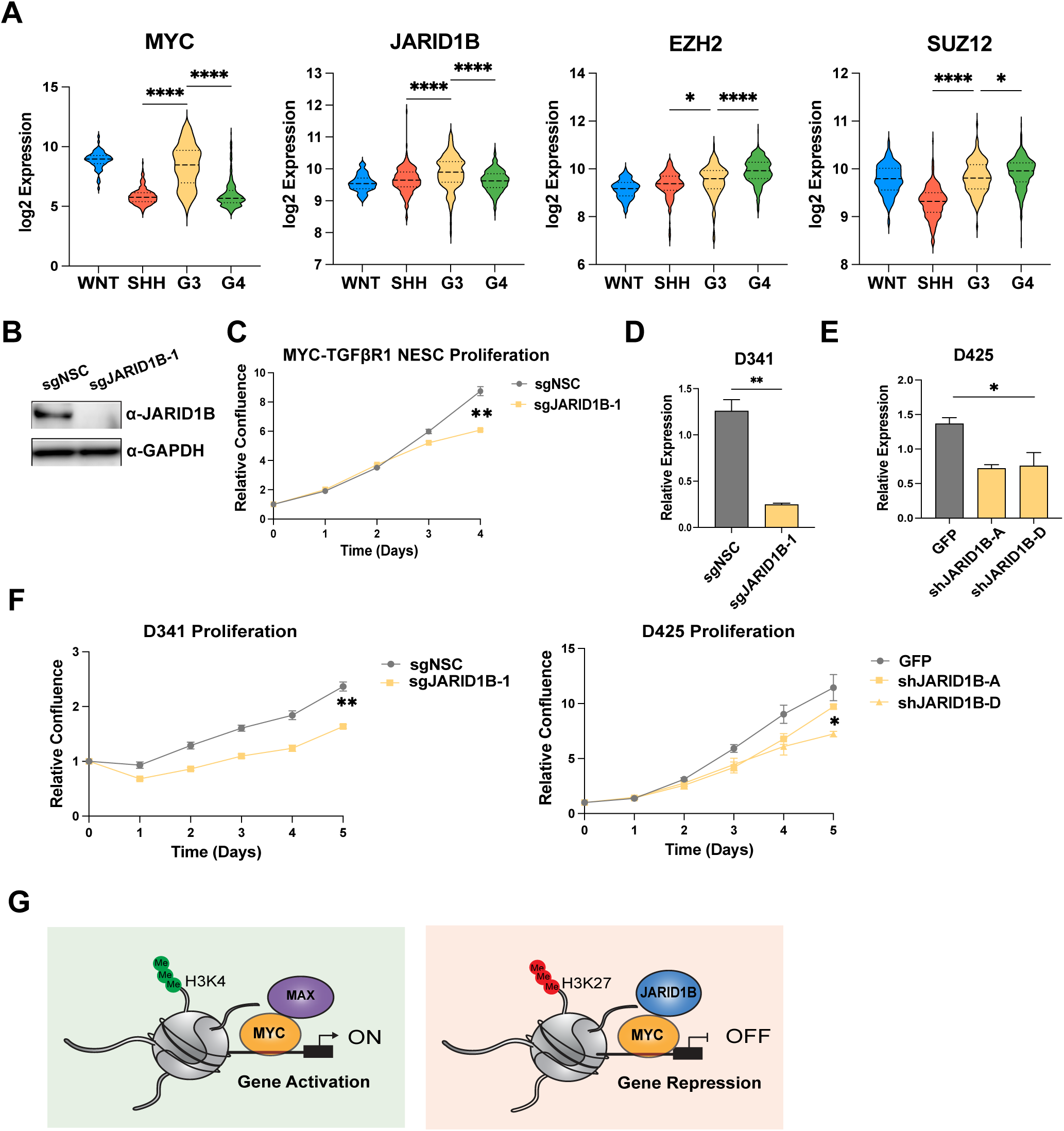
JARID1B is a potential therapeutic target for group 3 medulloblastoma. **A**, Boxplot of expression of PRC-interacting genes in MB subgroups from Cavalli et al., 2017. **B**, Immunoblot of JARID1B and GAPDH in MYC-TGFβR1 NESCs. **C**, Cell proliferation of MYC-TGFβR1control and JARID1B knockdown cells over 5 days. Percent confluence normalized to D0 for each line, average mean± SEM (n=3). **D**, **E,** qRT-PCR analysis for JARID1B in D341 and D425 G3MB cell lines. Relative expression normalized to GAPDH, mean± SEM (n=3) **F**, Cell proliferation of D341 and D425 control and JARID1B knockdown cells over 5 days, average mean± SEM (n=3). Statistical significance calculated using unpaired student’s t-test comparing control cells to knockdown, * p<0.05. **G**, Proposed mechanism for MYC-driven transcription in G3MB.

To assess JARID1B as a potential target in G3MB, we transduced MYC-TGFβR1 NESCs with ZIM3-KRABdCas9^46^ and knocked down JARD1B using CRISPRi (**Fig. 4B**). We observed significant ablation of growth over time in knockdown cells compared to control, which is specific to MYC overexpression (**Fig. 4C**). MYCN expressing NESCs, which we have shown to lead to transformation of SHHMB *in vivo*^29^, with reduced JARID1B expression were not affected (**Fig. S5B-C**), indicating that JARID1B was being recruited specifically by MYC to regulate neuronal gene expression. We further performed CRISPRi experiments in *MYC*-amplified human G3MB cell line D341 (**Fig. S3B**) for JARID1B and also observed striking decrease in proliferation in JARID1B knockdown cells (**Fig. 4D, F**). This was further supported by shRNA mediated knockdown of JARID1B in D425 (**Fig. 4E-F**), indicating a correlation of sensitivity between MYC expression and JARID1B dependency.

## Discussion

The lineage of origin plays a fundamental role in MB etiology and investigating the role of epigenetic and transcription factors in the correct developmental context will guide more effective therapies. Here we present new humanized isogenic models for G3MB driven by *MYC* amplification and overexpression of TGFβ pathway effectors ACVR2A, TGFβR1, and TGFβ1 that recapitulate the human tumor type. We functionally validate the cooperation of candidate driver genes in intrinsic drug resistance *in vitro* and tumor formation *in vivo*. To investigate mechanisms of MB tumor development, we queried MYC genomic localization along with gene expression profiling and observed that MYC is targeted to active chromatin as well as transcriptionally repressed regions regulated by JARID1B. Our findings further demonstrate that JARID1B has a critical role in promoting lineage plasticity and suppressing neurodevelopmental states in the context of MYC-driven G3MB (**Fig. 4G**). This supports JARID1B as a new therapeutic target specifically for MYC-driven G3MB for further exploration.

Targeting aberrantly altered epigenetic pathways has emerged as a promising strategy for cancer treatment with a recent focus in histone methylation^47^. Small inhibitors targeting JARID1B have been developed to suppress catalytic activity of the molecule in various disease models^48–52^. These small molecules are pan-inhibitors of the KDM5 family members, which contain the Jumonji-N domain (JmjN) and Jumonji-C domain (JmjC)^53–55^. JARID1B overexpression has been identified in several cancers and associated with poor outcomes^36,56^. Consistent with these findings, we also observe elevated levels of JARID1B in human G3MB. Moreover, overexpression of JARID1B causes dysregulation of neuronal differentiation during normal development^41,57^, suggesting these inhibitors could target the stem component of highly aggressive G3MB tumors. Outside of cancer, the first JmjC demethylase-based clinical trial studied the efficacy of the pan-KDM5 inhibitor GS-5801 in chronic hepatitis B^58^. While that study was unsuccessful, this highlights the need for therapies targeting H3K4 demethylation in cancer as well as other diseases and thus warrants more investigation.

The importance of JARID1B as a therapeutic target is further highlighted by biochemical studies supporting the protein as a potential binding partner for MYC. Secombe et al. demonstrated that the *Drosophila* ortholog for JARID1B, Lid, binds directly to dMyc to facilitate maintenance of H3K4me3 on the chromatin^55^. In mammalian systems, a ternary corepressor complex at promoters was identified containing MYC, TFAP2C, and JARID1B that downregulated the *CDKN1A* gene. Moreover, overexpression of all three proteins resulted in forced S-phase entry and attenuation of checkpoint activation and promoted chemotherapy resistance^59^. This suggests that MYC recruits JARID1B to dynamically silence neuronal development genes by removal of the H3K4me3 mark. We hypothesize that there is crosstalk that leads to MYC-mediated JARID1B recruitment to key neurodevelopmental promoters to maintain a blocked differentiation state.

Previous studies have reported that MYC promotes gene repression on the chromatin template and our data further supports a role for this mechanism in driving aggressive G3MB^60–62^. MYC is a master transcriptional regulator that binds open chromatin through interactions with EP300, TIP60, WDR5, as well as several other developmentally specific transcription factors^63^. In mouse models, repression by Myc-Miz1 suppresses both apoptosis and differentiation in G3MB^62^. The MYC-MIZ-1 interaction accounts for ∼40% of MYC-repressed genes^62^, suggesting additional interactors contribute to transcriptional repression. Notably, direct interaction of MYC with SMAD2/3 has been implicated to repress inhibitory functions of SMAD proteins and anti-proliferative functions of TGFβ signaling^64^. Tu et al. probed the MYC interactome and identified the histone methyltransferase G9a as a direct interactor in breast cancer, which was required for gene repression^61^. G9a and PRC2 have been shown to be physically interact in ESCs and bind a common set of developmental genes^65^. In a complementary manner, PRC2 recruits JARID1B to the DNA template and JARID1B is critical in regulating neural differentiation in ESCs^41^. Our data supports that JARID1B may be a context-specific binding partner to regulate MYC-mediated gene repression in MB in a similar manner to MYC-G9a interactions. This underscores the importance of finding new non-canonical interactors of MYC to better understand MYC-regulated transcription, which will give us a better understanding of how to target MYC-driven cancers.

In summary, we present new hiPSC models for G3MB that incorporate the developmental context that this tumors arises from and serve as an effective platform to understand key drivers of tumor formation, maintenance, and metastasis. Utilizing this model, we observe gains of H3K27me3 at neuronal differentiation genes that are regulated by MYC and JARID1B. We validate JARID1B as a new potential therapeutic target for G3MB and an important regulator of the chromatin landscape to promote a suppressed differentiated state. We demonstrate the potential to model different MB subtypes with specific genetic alterations that are not available in current cell lines. This has important significance for designing future targeted therapies for MB.

## Materials and Methods

### Experimental Model and Subject Details

#### Human iPSC Culture and differentiation to NESC

Human NESCs were generated from established human WTC10 iPSCs (gift from Bruce R. Conklin, Gladstone Institutes, UCSF) as previously described^27,29^. All experiments using iPSC in the Weiss lab (UCSF) were approved by Human GESCR Committee of the UCSF Stem Cell Research Oversight Committee. NESCs were cultured on poly-L-ornithine/laminin-coated wells in NESC media (DMEM/F-12 with GlutaMAX, 1X penicillin/streptomycin, 1X N2 supplement, 0.05X B27 without vitamin A supplement, 1.6 g/L Glucose, 20 ng/mL EGF, 20 ng/mL FGF2). NESCs passaged every three to four days using TrypLE Express and NESC media with 2uM of ROCK inhibitor Thiazovivin (TZV). Both hiPSC and NESCs were periodically karyotyped with Thermofisher KaryoStat assays.

#### G3MB Cell lines

D283, D341, and D425 cell lines were derived at Duke University and gifted from Dr. Martine Roussel’s lab (St. Jude’s). D283 were cultured in Eagles minimum media with 10% FBS and 1X penicillin/streptomycin in ultra-low flasks. D341 were cultured in Eagles minimum media with 20% FBS, 1X penicillin/streptomycin, and 1X Glutamax in ultra-low flasks. D425 were cultured in Neurobasal-A media supplemented with 1X B-27, 1X penicillin/streptomycin, 1X Glutamax, 20 ng/mL human recombinant EGF, 20 ng/mL human recombinant FGF, and 2 ug/mL Heparin solution in normal tissue culture flasks. Cell lines were periodically authenticated by STR genotyping (IDEXX BioAnalytics).

#### Animals

Immunocompromised NOD-*scid* IL2Rgamma^null^, NOD-*scid* IL2Rg^null^ (NSG) 6–8-week-old female mice used for transplantation were purchased from Jackson Labs or obtained from the UCSF Laboratory Animal Resource Center (LARC) Breeding Services. Mice were maintained in the LARC Animal Facility at UCSF. All experiments were performed in accordance with national guidelines and regulations, and with the approval of the IACUC and IBC at UCSF. For each *in vivo* experiment, 300,000 NESCs were resuspended in NESC media + TZV and injected orthotopically using a stereotactic frame into the cerebellum starting from lambda: 2 mm lateral right, 2 mm posterior, and 2 mm deep or into the lateral ventricle starting from bregma 1.5 mm lateral right, 3 mm posterior, 2 mm deep. Bioluminescence was measured in NSG mice injected with mScarlet-Luciferase or GFP-Luciferase tagged cells as previously described. Measurements were taken weekly 7 days post injection until mice reached endpoint. Mice were euthanized at endpoint, which was either signs of tumor growth (e.g. hunched back, weight loss, head tilt, neurological defects, etc.) or one year after transplantation.

### Method Details

#### Plasmids, Lentiviral Production, and Cell Transduction

HEK293T cells were grown in DMEM media supplemented with 10% FBS and 1X GlutaMAX. For lentiviral production, plasmids containing constructs of interest were co-transfected into HEK293T cells with the packaging plasmids. For each 10cm^2^ plate, the following DNAs were diluted in 1 mL Opti-MEM: 6μg construct, 4μg psPAX2, and 1μg pCMV-VSV-G. The DNA was combined with Trans-IT-Lenti Transfection reagent (Mirus) and incubated at room temperature for 10 minutes. The transfection mixture was then added dropwise onto the HEK293T cells with 10 mL of fresh media. The next day, media was changed to DMEM media supplemented with 30% FBS, 1X GlutaMAX and 1X Viralboost reagent (Alstem). Viral particles were harvested both at 48 hours and 72 hours after transfection by passing the cell culture medium through a 0.45μM PVDF syringe filter and supernatant was concentrated using Lenti-X (Takara) following manufacturer’s protocol. The pellet was resuspended to 1/10 of the original volume using PBS and used for transduction of recipient cultures or stored at -80°C.

#### RNA extraction, cDNA synthesis, and qRT-PCR

RNA was extracted using the Quick-RNA Miniprep kit (Zymo Research). 1 ug of RNA was converted to cDNA using the High-Capacity cDNA Reverse Transcription Kit (Applied Biosystems) and the following settings: 25°C for 10 min, 37°C for 120 min, and 85°C for 5 min. qPCR was performed using PowerUp SYBR green (Applied Biosystems) with the following settings: 95°C for 2 min, 40 cycles (95 °C for 15 s and 60 °C for 1 min). Primer sequences are listed in Table SX.

#### Immunoblotting

Cells were harvested following standard cell splitting procedures. Pellets were lysed in 200–300 µL Protein Lysis Buffer (PLB), depending on pellet size. Lysates were transferred to glass sonication tubes, sonicated for 5 cycles using the Bioruptor (Diagenode), and incubated on ice for 5 minutes. Samples were centrifuged at maximum speed for 20 minutes at 4°C. Protein was quantified using the BCA assay (ThermoFisher) and samples were prepared by mixing appropriate volumes of PLB, 4X sample buffer, and 10X reducing agent to achieve 30 µg of total protein per sample. Samples were heated at 70°C for 10 minutes and briefly vortexed, centrifuged, and loaded onto precast Bolt Bis-Tris gels (Invitrogen). Proteins were separated and transferred to PVDF membrane (0.45μM pore size). Membranes were blocked in 5% milk in TBST for 1 hour at room temperature and incubated with primary antibodies diluted in 5% milk or BSA in TBST overnight at 4°C. The next day, the membranes were washed 3X in TBST, incubated with secondary antibodies diluted in 5% milk in TBST for 2 hours at room temperature, and followed with 3 additional TBST washes. The membranes were imaged with ECL chemiluminescence reagent (Biorad) and visualized using the ImageQuant 800 Western blot imaging system (Amersham). If necessary, membranes were stripped using stripping buffer for 10 minutes at room temperature, washed with DI water, and re-blocked before re-probing.

#### Cell proliferation assays

Equal number of NESCs were plated into 24- or 96-well plates and treated with TGFβR1 inhibitors Galunisertib and LY3200882 (Selleck Chemicals) and chemotherapeutic agents cyclophosphamide (Santacruz) and carboplatin (Selleck Chemicals). Media was changed every other days with fresh inhibitor for up to 5 days. Cells were imaged and quantified using the Incucyte™ Live-Cell Imaging System every 12-24 hours during treatment.

#### RNA-seq

Tumor tissue was homogeneous using a QIAshredder (Qiagen) and cell and tissue RNA was extracted using the Quick-RNA extraction kit (Zymo Research). RNA samples were quantified on the Qubit and sent to Signios Bio for library preparation using the Illumina Stranded mRNA kit and sequenced on the Novaseq X targeting 80 million paired end 100 bp reads per sample.

#### CUT&RUN

Approximately 500,000 cells per sample for histone modifications, FLAG, and MYC were harvested for CUT&RUN. CUT&RUN was performed using the CUTANA® ChIC/CUT&RUN Kit (EpiCypher, Cat#14–1048) according to the manufacturer’s instructions. Briefly, cells were incubated concanavalin A conjugated paramagnetic beads. The cell-bead solution was incubated with each respective antibody per sample overnight at 4°C. The next day, samples were incubated with Protein A-Protein G Micrococcal Nuclease (pAG-MNase). Following binding, the pAG-MNase was activated by the addition of CaCl_2_ and incubated for 2 hours. DNA was subsequently purified using SPRIselect Bead-Based Reagent (Beckman Coulter, Inc.) and eluted using TE Buffer. DNA was then quantified Qubit fluorometer and libraries were prepared using the CUTANA™ CUT&RUN Library Prep Kit (EpiCypher 14-1001 / 14-1002). Libraries were sequenced at the UCSF Center for Advanced Technology (CAT) on an Illumina Novaseq X targeting 10 million 50 bp paired end reads per sample.

### Quantification and Statistical Analysis

#### RNA-seq data processing

Sequencing reads for samples were aligned to the reference genome GRCh38/hg38 by STAR v2.7.11a^66^. Read counts per gene were tallied by featureCounts^67^ from the R package, Rsubread v2.16.1, using Gencode v45 reference genome. Genes with fewer than 10 counts total amongst the sample groups were excluded from downstream analysis. Differential gene expression analysis was performed with DESeq2 v1.42.1^68^ which was used to calculate log2FC and Benjamini-Hochberg adjusted p-values between sample groups. The cut off for significant differentially expressed genes was an adjusted p-value < 0.05 and L2FC > |1|. Heatmap generation was completed with DESeq2 variance stabilized counts and ComplexHeatmap v2.18.0^69^. Volcano plots were made with EnhancedVolcano v1.20.0 in RStudio^70^

#### ChEA and Pathway Enrichment Analysis

GSEA, Gene Ontology (GO) and ChEA analysis was performed as previously described^71^ using clusterProfiler^33^ and Enrichr^72^. Plots were visualized using RStudio v4.3.1^70^

#### Classification of NESC-derived tumors

Raw RNA-seq reads were processed to remove low-quality sequences and adapter sequences. To discriminate between human and mouse reads, *disambiguate* was used to filter out reads with a higher mapping quality score in the mouse genome compared to the human genome^73^. Human-specific reads were mapped to GRCh38/hg38 using STAR to generate *Fragments Per Kilobase of transcript per Million* mapped reads (FPKM) values. To quantify the resemblance of NES-derived tumors with medulloblastoma subgroups, we trained a machinelearning classification model on a large series of primary medulloblastoma tumors. Specifically, two published gene expression microarray series of medulloblastomas, with medulloblastoma subgroup determined by DNA methylation, were obtained (GEO: GSE85218 and EGAS00001001953). After removing duplicate samples this resulted in 1112 unique expression profiles that underwent Robust Multi-array Average normalization. Because gene sets are more robust to platform differences than individual genes, single-sample gene set enrichment (ssGSEA) scores were derived by using all *MSigDB* gene sets that contained ≥ 80% mouse–human orthologs and ≥ 15 and ≤ 400 genes^74^. A random forest classification model was then trained using a 70–30 training-test split of the medulloblastoma expression profiles, using the gene set enrichment scores as features to predict medulloblastoma subgroup. Gene sets with a scaled feature importance score > 8 were used to build the final classification model resulting in a total of 409 features (test set accuracy = 0.956). This model was used to predict the subgroup class probabilities for all NES-derived tumors. The same workflow was utilized to create a binary machine learning model to classify if tumors were medulloblastoma. The dataset used to train the non-medulloblastoma model consisted of 1473 primary CNS tumors, after filtering undefined entities, encompassing 10 distinct CNS tumor types obtained from the R2 pediatric genome portal (accession “Tumor Brain (DKFZ-public)- Kool-1678-MAS5.0-u133p2”). Using a 70–30 training-test split, the non-medulloblastoma model achieved a test set accuracy of 0.987.

#### Analysis of RNA expression data

Publicly available RNA expression and relevant clinical prognostic parameters were obtained from the R2: Genomics Analysis and Visualization Platform (https://hgserver1.amc.nl/cgi-bin/r2/main.cgi) from following datasets: Cavalli and MAGIC.

#### CUT&RUN data processing

Sequenced reads from CUT&RUN experiments were analyzed as previously described^75^. trimmed for adapter sequences using TrimGalore with default parameters. Reads were aligned to reference genome hg38 using bowtie2 with parameter (–maxins 2000) for paired-end samples. The aligned reads were filtered for alignment quality q30 and sorted using Samtools v1.9 (default parameters) followed by duplicate removal using Picard (default parameters). Peak calling was performed using Macs2 callpeak commands. The input-corrected signal tracks (bigwig) were obtained using Macs2 bdgcmp command with parameters (–method FE), followed by bedClip and bedGraphToBigWig commands from UCSC Genome Browser Tools^76^, both with default parameters. QC statistics are reported in Supplementary Data.

#### Genome Distribution Analysis

To generate genome distribution analysis plots, MACS2-called peaks were merged with SAMtools^77^ to find the union between sample replicates. Merged sample peaks were annotated via ChIPseeker v1.38.0^78^ with UCSC hg38. Differential binding analysis was completed using DiffBind v3.12.0^39^ with its inbuilt DESeq2 method to call significance between samples. Motif analysis was completed with the MEME Suite MEME-ChIP^79^.

#### Metagene, Heatmaps, and Boxplots

Analysis was performed as previously described^71^. Metagene, heatmaps and boxplots of enrichment at promoters or genes were generated with deepTools and R Studio. Heatmaps and boxplots were generated using R Studio.

#### Statistical Analysis

All statistical analyses were performed using GraphPad Prism10. Differences in means were determined with unpaired Student’s t-test or ANOVA. Differences in survival were determined using the log-rank test. In general, statistical significance is shown using asterisks (*p<0.05, **p<0.01, ***p<0.0001), but exact p*-*values are provided in figure panels or figure legends whenever possible. Statistical tests performed as part of the RNA-seq and CUT&RUN analyses are detailed in relevant sections above.

#### Data availability

All sequencing data will be submitted to NCBI Gene Expression Omnibus at the time of submission.

## Acknowledgements

The authors thank David R. Raleigh, S. John Liu as well as current and former members of the Weiss lab for advice, reagents, and technical support. We thank Martine Roussel and her lab for sharing human G3MB cell lines as well as support and advice. We also thank Luke Gilbert for providing CRISPR-Cas9 vectors. We also acknowledge the support of Anny Shai and the staff of the UCSF Brain Tumor Center Biorepository and Pathology Core (NIH P50 CA097257), UCSF Helen Diller Family Comprehensive Cancer Center LCA and LCA-Genomic Core Facility (NIH P30CA082103), Eric Chow and the staff of the UCSF Center for Advanced Technology supported by UCSF PBBR, RRP IMIA, and NIH 1S10OD028511-01 grants. These studies were supported by the NIH grant T32 CA151022, Damon Runyon-Sohn Pediatric Fellowship Award, Alex Lemonade Stand Foundation Young Investigator Grant, and AACR-Sontag Foundation Brain Cancer Research Fellowship (Z.A.Q);.

## Author Contributions

Z.A.Q., M.H., and W.A.W. conceived and designed the overall study. Z.A.Q established the NESC models and performed mouse experiments with E.H., D.S., S.W., L. W., and R.P. E.H., M.O.J., and S.G. performed proliferation and drug experiments. E.H. and D.S. performed histological experiments and slides were analyzed by E.G. Z.A.Q. performed RNA-seq and CUT&RUN experiments and analysis was performed by D.S. and W.M.I. B.G., K.S., and L.H. performed RNA-seq clustering analysis to human MB. Z.A.Q., E.H., and D.S. performed molecular biology experiments. Gene vectors and experimental advice was provided by M.H. A.G.M., P.A.N., and M.D.T. supervised computational analyses. Z.A.Q and W.A.W. wrote the manuscript with contributions from all authors.

## Conflict of Interest Statement

The authors declare that they have no competing interests related to this project.

## Notes

### Competing Interest Statement

The authors have declared no competing interest.

## References

1. Hovestadt, V. et al. Medulloblastomics revisited: biological and clinical insights from thousands of patients. Nature Reviews Cancer vol. 20 42–56 (2020).

2. Cacciotti, C., Fleming, A. & Ramaswamy, V. Advances in the molecular classification of pediatric brain tumors: a guide to the galaxy. Journal of Pathology vol. 251 249–261 (2020).

3. Van Ommeren, R., Garzia, L., Holgado, B. L., Ramaswamy, V. & Taylor, M. D. The molecular biology of medulloblastoma metastasis. Brain Pathol. 30, 691–702 (2020).

4. Northcott, P. A. et al. Medulloblastoma Comprises Four Distinct Molecular Variants. J. Clin. Oncol. 29, 1408–1414 (2011).

5. Taylor, M. D. et al. Molecular subgroups of medulloblastoma: the current consensus. Acta Neuropathol. 123, 465–472 (2012).

6. Hovestadt, V. et al. Decoding the regulatory landscape of medulloblastoma using DNA methylation sequencing. Nature 510, 537–541 (2014).

7. Juraschka, K. & Taylor, M. D. Medulloblastoma in the age of molecular subgroups: a review: JNSPG 75th Anniversary Invited Review Article. J. Neurosurg. Pediatr. 24, 353–363 (2019).

8. Northcott, P. A. et al. Subgroup-specific structural variation across 1,000 medulloblastoma genomes. Nature 488, 49–56 (2012).

9. Lin, C. Y. et al. Active medulloblastoma enhancers reveal subgroup-specific cellular origins. Nature 530, 57–62 (2016).

10. Akhurst, R. J. Targeting TGF-β Signaling for Therapeutic Gain. Cold Spring Harb. Perspect. Biol. 9, a022301 (2017).

11. Massagué, J. TGFβ signalling in context. Nat. Rev. Mol. Cell Biol. 13, 616–630 (2012).

12. Oshimori, N., Oristian, D. & Fuchs, E. TGF-β promotes heterogeneity and drug resistance in squamous cell carcinoma. Cell 160, 963–976 (2015).

13. Dawson, M. & Kouzarides, T. Cancer Epigenetics: From Mechanism to Therapy. Cell 150, 12–27 (2012).

14. Northcott, P. A. et al. Enhancer hijacking activates GFI1 family oncogenes in medulloblastoma. Nature 511, 428–434 (2014).

15. Roussel, M. F. & Stripay, J. L. Epigenetic Drivers in Pediatric Medulloblastoma. Cerebellum 17, 28–36 (2018).

16. Dawson, M. A. et al. Cancer epigenetics: from mechanism to therapy. Cell 150, 12–27 (2012).

17. Hendrikse, L. D. et al. Failure of human rhombic lip differentiation underlies medulloblastoma formation. Nat. 2022 6097929 609, 1021–1028 (2022).

18. Lee, J. J. Y. et al. ZIC1 is a context-dependent medulloblastoma driver in the rhombic lip. Nat. Genet. 2025 571 57, 88–102 (2025).

19. Smith, K. S. et al. Unified rhombic lip origins of group 3 and group 4 medulloblastoma. Nature 609, 1012–1020 (2022).

20. Strahl, B. D. & Allis, C. D. The language of covalent histone modifications. Nature 403, 41–45 (2000).

21. Piunti, A. & Shilatifard, A. The roles of Polycomb repressive complexes in mammalian development and cancer. Nature Reviews Molecular Cell Biology vol. 22 (2021).

22. Cenik, B. K. & Shilatifard, A. COMPASS and SWI/SNF complexes in development and disease. Nature Reviews Genetics vol. 22 (2021).

23. Sanghrajka, R. M. et al. KMT2D suppresses Sonic hedgehog-driven medulloblastoma progression and metastasis. iScience 26, (2023).

24. Haldipur, P. et al. Spatiotemporal expansion of primary progenitor zones in the developing human cerebellum. Science 366, 454–460 (2019).

25. Aldinger, K. A. et al. Spatial and cell type transcriptional landscape of human cerebellar development. Nat. Neurosci. 24, 1163–1175 (2021).

26. Vladoiu, M. C. et al. Childhood cerebellar tumours mirror conserved fetal transcriptional programs. Nature 572, 67–73 (2019).

27. Tailor, J. et al. Stem Cells Expanded from the Human Embryonic Hindbrain Stably Retain Regional Specification and High Neurogenic Potency. J. Neurosci. 33, 12407–12422 (2013).

28. Koch, P., Opitz, T., Steinbeck, J. A., Ladewig, J. & Brüstle, O. A rosette-type, self-renewing human ES cell-derived neural stem cell with potential for in vitro instruction and synaptic integration. Proc. Natl. Acad. Sci. U. S. A. 106, 3225–30 (2009).

29. Huang, M. et al. Engineering Genetic Predisposition in Human Neuroepithelial Stem Cells Recapitulates Medulloblastoma Tumorigenesis. Cell Stem Cell 25, 433–446.e7 (2019).

30. Donati, G. & Amati, B. MYC and therapy resistance in cancer: risks and opportunities. Molecular Oncology vol. 16 (2022).

31. Teixeira, A. F., ten Dijke, P. & Zhu, H.-J. On-Target Anti-TGF-β Therapies Are Not Succeeding in Clinical Cancer Treatments: What Are Remaining Challenges? Front. Cell Dev. Biol. 0, 605 (2020).

32. Jia, S., Wu, D., Xing, C. & Meng, A. Smad2/3 activities are required for induction and patterning of the neuroectoderm in zebrafish. Dev. Biol. 333, 273–284 (2009).

33. Yu, G., Wang, L. G., Han, Y. & He, Q. Y. ClusterProfiler: An R package for comparing biological themes among gene clusters. Omi. A J. Integr. Biol. 16, (2012).

34. Lachmann, A. et al. ChEA: transcription factor regulation inferred from integrating genome-wide ChIP-X experiments. Bioinformatics 26, 2438–44 (2010).

35. Ren, Z. et al. A PRC2-Kdm5b axis sustains tumorigenicity of acute myeloid leukemia. Proc. Natl. Acad. Sci. U. S. A. 119, (2022).

36. Hinohara, K. et al. KDM5 Histone Demethylase Activity Links Cellular Transcriptomic Heterogeneity to Therapeutic Resistance. Cancer Cell 34, (2018).

37. Wu, T. et al. clusterProfiler 4.0: A universal enrichment tool for interpreting omics data. Innovation 2, (2021).

38. Hammal, F., De Langen, P., Bergon, A., Lopez, F. & Ballester, B. ReMap 2022: a database of Human, Mouse, Drosophila and Arabidopsis regulatory regions from an integrative analysis of DNA-binding sequencing experiments. Nucleic Acids Res. 50, D316–D325 (2022).

39. Stark, R. & Brown, G. DiffBind : differential binding analysis of ChIP-Seq peak data. Bioconductor (2011).

40. Albert, M. et al. The Histone Demethylase Jarid1b Ensures Faithful Mouse Development by Protecting Developmental Genes from Aberrant H3K4me3. PLoS Genet. 9, (2013).

41. Schmitz, S. U. et al. Jarid1b targets genes regulating development and is involved in neural differentiation. EMBO J. 30, (2011).

42. Hayami, S. et al. Overexpression of the JmjC histone demethylase KDM5B in human carcinogenesis: Involvement in the proliferation of cancer cells through the E2F/RB pathway. Mol. Cancer 9, (2010).

43. Roesch, A. et al. A Temporarily Distinct Subpopulation of Slow-Cycling Melanoma Cells Is Required for Continuous Tumor Growth. Cell 141, (2010).

44. Xiang, Y. et al. JARID1B is a histone H3 lysine 4 demethylase up-regulated in prostate cancer. Proc. Natl. Acad. Sci. U. S. A. 104, (2007).

45. Yamamoto, S. et al. JARID1B is a luminal lineage-driving oncogene in breast cancer. Cancer Cell 25, (2014).

46. Replogle, J. M. et al. Maximizing CRISPRi efficacy and accessibility with dual-sgRNA libraries and optimal effectors. Elife 11, (2022).

47. Gray, Z. H., Honer, M. A., Ghatalia, P., Shi, Y. & Whetstine, J. R. 20 years of histone lysine demethylases: From discovery to the clinic and beyond. Cell 188, 1747–1783 (2025).

48. Tumber, A. et al. Potent and Selective KDM5 Inhibitor Stops Cellular Demethylation of H3K4me3 at Transcription Start Sites and Proliferation of MM1S Myeloma Cells. Cell Chem. Biol. 24, (2017).

49. Thomas, L. R. et al. Interaction of the oncoprotein transcription factor MYC with its chromatin cofactor WDR5 is essential for tumor maintenance. Proc. Natl. Acad. Sci. U. S. A. 116, 25260–25268 (2019).

50. Johansson, C. et al. Structural analysis of human KDM5B guides histone demethylase inhibitor development. Nat. Chem. Biol. 12, (2016).

51. Vinogradova, M. et al. An inhibitor of KDM5 demethylases reduces survival of drug-tolerant cancer cells. Nat. Chem. Biol. 12, (2016).

52. Paroni, G. et al. HER2-positive breast-cancer cell lines are sensitive to KDM5 inhibition: definition of a gene-expression model for the selection of sensitive cases. Oncogene 38, (2019).

53. Klose, R. J. et al. The Retinoblastoma Binding Protein RBP2 Is an H3K4 Demethylase. Cell 128, (2007).

54. Christensen, J. et al. RBP2 Belongs to a Family of Demethylases, Specific for Tri-and Dimethylated Lysine 4 on Histone 3. Cell 128, (2007).

55. Secombe, J., Li, L., Carlos, L. & Eisenman, R. N. The Trithorax group protein Lid is a trimethyl histone H3K4 demethylase required for dMyc-induced cell growth. Genes Dev. 21, (2007).

56. Jose, A. et al. Histone demethylase KDM5B as a therapeutic target for cancer therapy. Cancers vol. 12 (2020).

57. Dey, B. K. et al. The Histone Demethylase KDM5b/JARID1b Plays a Role in Cell Fate Decisions by Blocking Terminal Differentiation. Mol. Cell. Biol. 28, (2008).

58. Phillips, S., Jagatia, R. & Chokshi, S. Novel therapeutic strategies for chronic hepatitis B. Virulence vol. 13 (2022).

59. Wong, P.-P. et al. Histone Demethylase KDM5B Collaborates with TFAP2C and Myc To Repress the Cell Cycle Inhibitor p21 cip ( CDKN1A ). Mol. Cell. Biol. 32, (2012).

60. Caputo, V. S. et al. Myc and Bet Proteins Orchestrate the Early Regulatory Genome Changes Required for Osteoclast Lineage Commitment. Blood 134, (2019).

61. Tu, W. B. et al. MYC Interacts with the G9a Histone Methyltransferase to Drive Transcriptional Repression and Tumorigenesis. Cancer Cell 34, (2018).

62. Vo, B. H. T. et al. The Interaction of Myc with Miz1 Defines Medulloblastoma Subgroup Identity. Cancer Cell 29, 5–16 (2016).

63. Kress, T. R., Sabò, A. & Amati, B. MYC: Connecting selective transcriptional control to global RNA production. Nature Reviews Cancer vol. 15 (2015).

64. Feng, X. H., Liang, Y. Y., Liang, M., Zhai, W. & Lin, X. Direct interaction of c-Myc with Smad2 and Smad3 to inhibit TGF-β-mediated induction of the CDK inhibitor p15Ink4B. Mol. Cell 9, (2002).

65. Mozzetta, C. et al. The Histone H3 Lysine 9 Methyltransferases G9a and GLP Regulate Polycomb Repressive Complex 2-Mediated Gene Silencing. Mol. Cell 53, (2014).

66. Dobin, A. et al. STAR: ultrafast universal RNA-seq aligner. Bioinformatics 29, 15–21 (2013).

67. Liao, Y., Smyth, G. K. & Shi, W. featureCounts: an efficient general purpose program for assigning sequence reads to genomic features. Bioinformatics 30, 923–930 (2014).

68. Love, M. I., Huber, W. & Anders, S. Moderated estimation of fold change and dispersion for RNA-seq data with DESeq2. Genome Biol. 15, 550 (2014).

69. Gu, Z., Eils, R. & Schlesner, M. Complex heatmaps reveal patterns and correlations in multidimensional genomic data. Bioinformatics 32, (2016).

70. RStudio Team. RStudio: Integrated Development for R. (2022).

71. Qadeer, Z. A. et al. ATRX In-Frame Fusion Neuroblastoma Is Sensitive to EZH2 Inhibition via Modulation of Neuronal Gene Signatures. Cancer Cell 36, (2019).

72. Chen, E. Y. et al. Enrichr: interactive and collaborative HTML5 gene list enrichment analysis tool. BMC Bioinformatics 14, 128 (2013).

73. Ahdesmäki, M. J., Gray, S. R., Johnson, J. H. & Lai, Z. Disambiguate: An open-source application for disambiguating two species in next generation sequencing data from grafted samples. F1000Research 5, (2017).

74. Subramanian, A. et al. Gene set enrichment analysis: a knowledge-based approach for interpreting genome-wide expression profiles. Proc. Natl. Acad. Sci. U. S. A. 102, 15545–50 (2005).

75. Mohammed Ismail, W., et al. MacroH2A histone variants modulate enhancer activity to repress oncogenic programs and cellular reprogramming. *Commun*. Biol. 6, (2023).

76. Kent, W. J., Zweig, A. S., Barber, G., Hinrichs, A. S. & Karolchik, D. BigWig and BigBed: Enabling browsing of large distributed datasets. Bioinformatics 26, (2010).

77. Li, H. et al. The Sequence Alignment/Map format and SAMtools. Bioinformatics 25, (2009).

78. Yu, G., Wang, L. G. & He, Q. Y. ChIP seeker: An R/Bioconductor package for ChIP peak annotation, comparison and visualization. Bioinformatics 31, (2015).

79. Machanick, P. & Bailey, T. L. MEME-ChIP: Motif analysis of large DNA datasets. Bioinformatics 27, (2011).

