## Supplemental Figure 1 for "Human stem cell models for group 3 medulloblastoma uncover JARID1B as a regulator of the chromatin landscape"

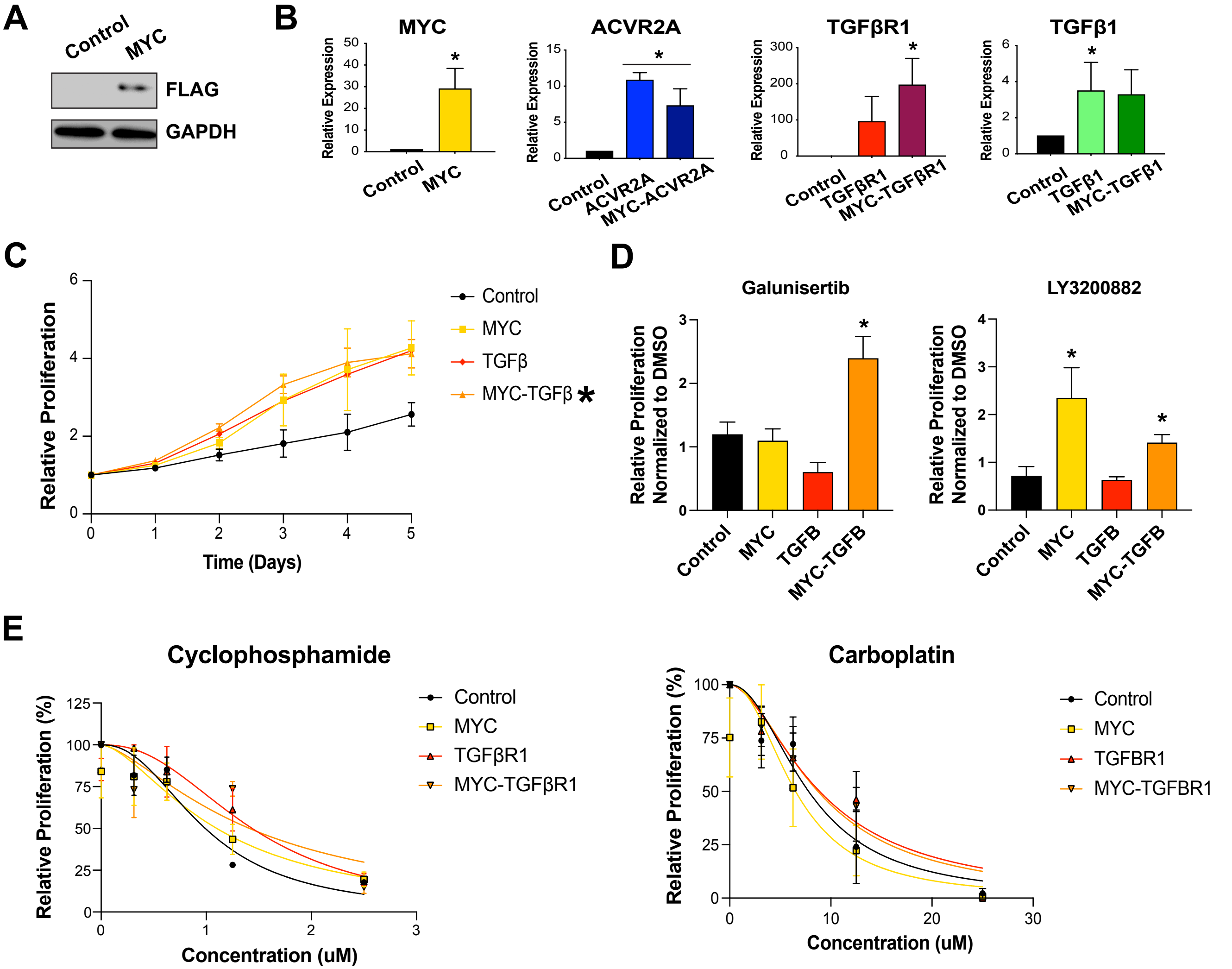


**Figure S1. Generation of NESCs expressing MYC and TGFβ. A**, Immunoblot for MYC and GAPDH in NESCs. **B**, qRT-PCR analysis in control, MYC and TGFβ expressing NESCs. Relative expression normalized to GAPDH, mean n± SEM (n=3). **C**, Proliferation over time (5 days) of NESCs expressing TGFβ effectors and/or MYC. Percent confluence normalized to D0 for each line, average mean± SEM (n=3). Statistical analysis performed using student unpaired t-test compared to control NESCs, p values represented as * and denotes p value<0.05. **D**, Proliferation of NESCs expressing TGFβ effectors ± MYC as indicated treated with 5 uM of TGFβRIi Galunisertib and LY3200882 for 5 days. Data normalized to DMSO for each cell line, average mean± SEM (n= 3). Statistical analysis performed using one-way ANOVA and individual comparisons assessed with the Benjamini-Hochberg proecudre, p values represented as * and denotes p value<0.05. **E**, Proliferation of NESCs expressing TGFβRI ± MYC as indicated treated with increasing concentration of drug for 4 days. Percent confluence normalized between 0 and 100% for the lowest and highest number for each cell line, average mean± SEM (n=3).
