## Supplemental Figure 2 for "Human stem cell models for group 3 medulloblastoma uncover JARID1B as a regulator of the chromatin landscape"

Qadeer et al., Figure S2

A

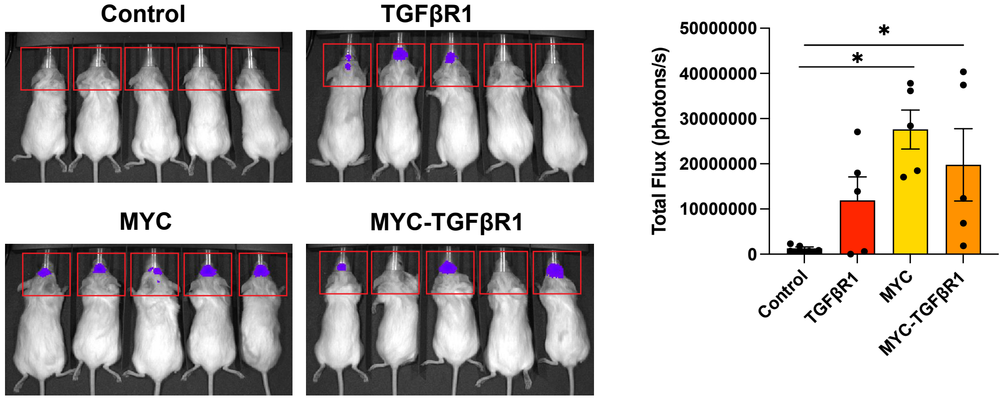


B

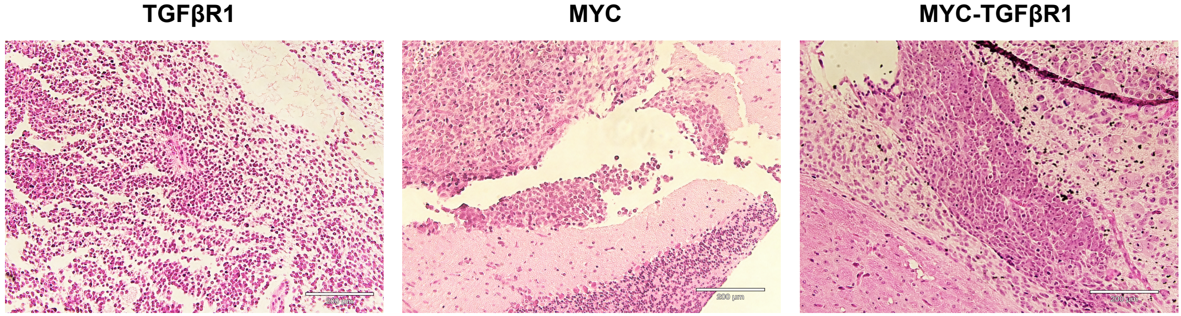


**Figure S2. Characterization of NESCs-derived tumors expressing MYC and TGFβR1. A**, Representative BLI images demonstrating aggressive tumor growth at D28 for implanted MYC alone and MYC+TGFβR1 NESCs. Statistical signifcanc calculated using unpaired Student's t-test, * denotes p-value <0.05 compared to mice implanted with control NESCs. **B**, H&E staining of tumors derived from TGFβR1 alone, MYC alone, and MYC+TGFβR1 expressing NESCs show features representative of MB. Images taken on the Echo Revolve at 10X mag.
