## Supplemental Figure 3 for "Human stem cell models for group 3 medulloblastoma uncover JARID1B as a regulator of the chromatin landscape"

### Qadeer et al., Figure S3

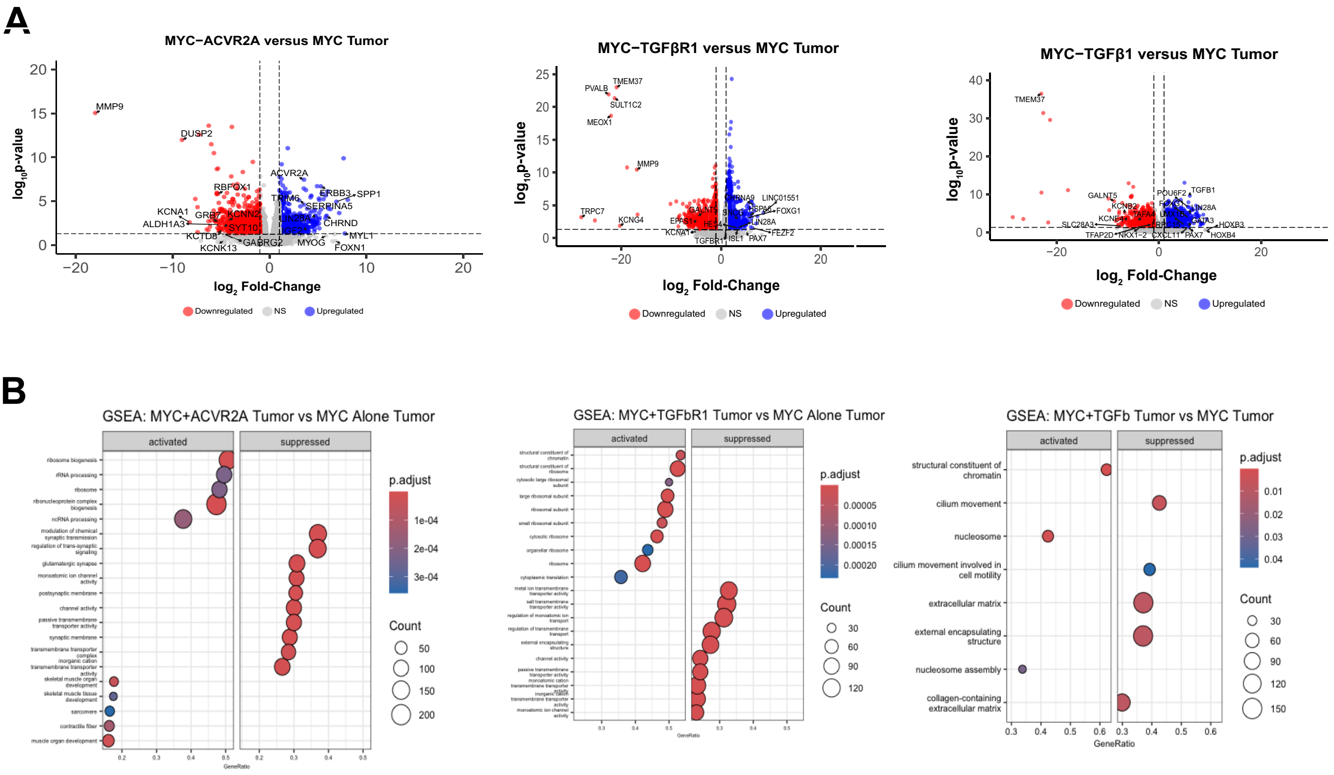


**Figure S3. RNA-seq analysis of MYC-TGFβ driven NESC-derived tumors. A**, Volcano plots comparing differentially expressed genes (DEG) between NESC-derived tumors driven by MYC-TGFβ versus MYC alone. Significant DEGs were called with log2FC ≥1 for Upregulated and log2FC ≤1 and Downregulated, p<0.05. **B**, Gene set enrichment analysis (GSEA) of significant DEGs for each genotype using clusterProfiler.
