## Supplemental Figure 4 for "Human stem cell models for group 3 medulloblastoma uncover JARID1B as a regulator of the chromatin landscape"

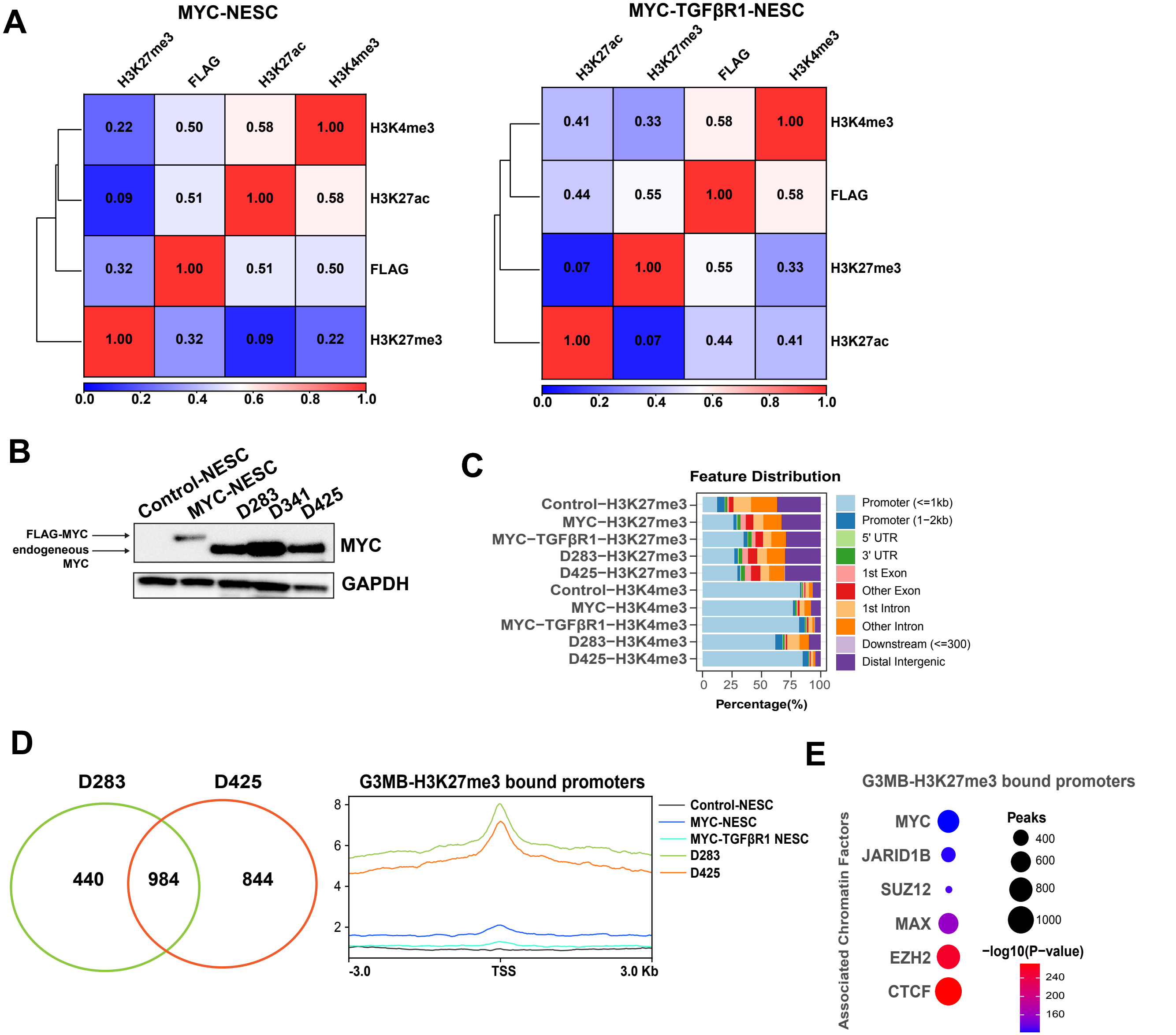


**Figure S4. CUT&RUN analysis of MYC-TGFβR1 driven NESCs and human G3MB cell lines. A**, Heatmap showing Spearman correlation of CUT&RUN signal for histone marks and FLAG-MYC for each line with FLAG peaks correlating more with H3K27me3 in MYC-TGFβR1 NESCs. **B**, Immunoblot analysis for MYC and GAPDH expression in NESCs G3MB cell lines. **C**, Genome wide distribution of significant peaks called for respective histone PTMs in NESCs and G3MB cell lines. **D**, Venn diagram (left) comparing differential D283/D425 H3K27me3-bound promoters gained over control-NESCs. Metagene profiles (right) of H3K27me3 at differentially bound promoters in MYC-driven G3MB cell lines. **E**, RemapEnrich analysis showing these genes are MYC and JARID1B targets.
