## Supplemental Figure 5 for "Human stem cell models for group 3 medulloblastoma uncover JARID1B as a regulator of the chromatin landscape"

Qadeer et al., Figure S5

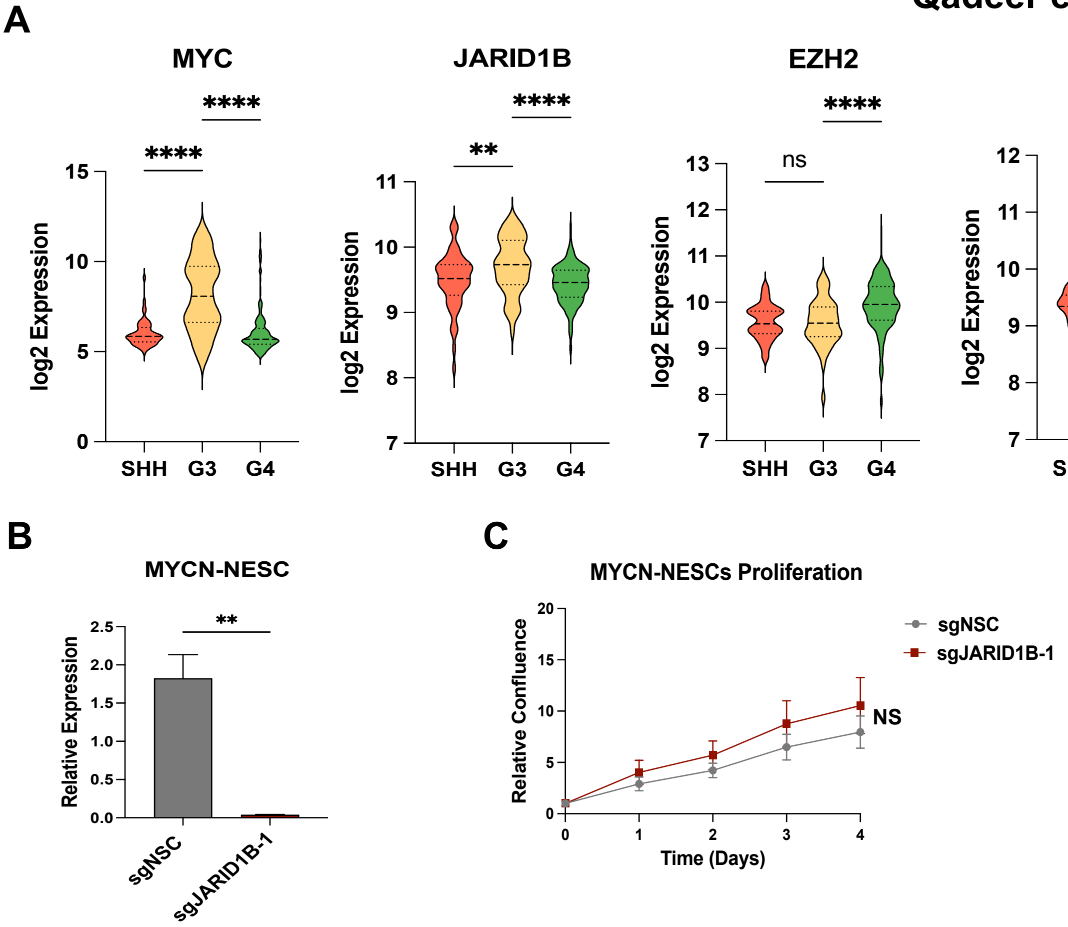


**Figure S5. JARID1B analysis in human MB and NESC cell lines. A**, Boxplot of expression of PRC-interacting genes in MB subgroups in the MAGIC dataset. **B**, qRT-PCR analysis for JARID1B in MYCN-expressing NESCs. Relative expression normalized to GAPDH, mean± SEM (n=3) **C**, Cell proliferation of MYCN-NESC control and JARID1B knockdown cells over 5 days. Percent confluence normalized to D0 for each line, average mean± SEM (n=3). Statistical signifance calculated using unpaired student's t-test or one-way ANOVA with Tukey's multiple comparision test. (*, p<0.05, **, p<0.01; ***, p<0.001, ****, p<0.0001)
